# GOA-1 regulates spermathecal transits

**DOI:** 10.1101/2025.09.30.679514

**Authors:** Fereshteh Sadeghian, Virginie Sjoelund, Erin J Cram

## Abstract

G protein signaling regulates calcium dynamics and contractility in the *C. elegans* spermatheca. G protein-coupled receptors activate heterotrimeric G proteins, triggering downstream cascades, including the Gαs-mediated activation of adenylyl cyclase and subsequent Protein Kinase A (PKA) activation. Our previous work identified GSA-1/Gαs and PKA as key modulators of Ca^2+^ oscillations and tissue contractility within the *C. elegans* spermatheca. In this study, we show that the inhibitory Gαi/o subunit GOA-1 regulates spermathecal transits. We employed TurboID proximity labeling and mass spectrometry to identify 16 candidate interactors of GOA-1. Depletion of these candidates by RNAi did not yield overt spermathecal transit defects.

## Description

Heterotrimeric G proteins are composed of an alpha (α), and a beta (β) gamma (γ) subunit, which, when activated by an upstream G protein-coupled receptor (GPCR) or G protein regulator (GPR), disassociate and independently trigger signaling cascades (Jastrzebska, 2013). In smooth muscle, the activation of Gαs initiates the activation of adenylyl cyclase (AC), which converts adenosine triphosphate (ATP) into 3’-5’-cyclic adenosine monophosphate (cAMP). Protein kinase A (PKA) becomes active when cAMP binds, leading to release of inhibition by the PKA regulatory subunits. PKA regulates various metabolic pathways (Lee et al., 2016), cell migration (Howe, 2004), and the relaxation of airway smooth muscle (Billington et al., 2013), among other functions (Sadeghian et al., 2022; Torres-Quesada et al., 2017).

The *C. elegans* spermatheca stores sperm and is the site of fertilization. In the spermatheca, acto-myosin contractility is stimulated by Ca^2+^ signaling via the phospholipase PLC-1, which stimulates the release of Ca^2+^ from the endoplasmic reticulum. In previous work, we identified GSA-1 (Gαs) signaling through PKA as an important regulator of coordinated Ca^2+^ signaling in the spermatheca (Castaneda et al., 2020). Oocyte entry initiates a series of Ca^2+^ oscillations, which lead to contraction and the exit of the fertilized egg into the uterus. Several crucial questions remain, such as what initiates this signaling cascade and maintains these Ca^2+^ oscillations and how the combined perception of mechanical and biochemical cues leads to proper response observed at the tissue level.

Here we show that, in addition to GSA-1 (Gαs), GOA-1 (Gi/o), an inhibitory G protein, also regulates spermathecal contractility. A *goa-1p*::GOA-1::GFP construct is expressed in the spermatheca and other tissues (Figure 1A). Depletion of GOA-1 by RNAi results in a failure of embryos to exit the spermatheca (Figure 1B). Worms expressing the Ca^2+^ sensor GCaMP under the *fln-1* promoter were subjected to control and *goa-1* RNAi. In this experiment, the percentage of spermathecae occupied with an oocyte was determined for each gene. If the gene is required for oocyte transit through the spermatheca, the percentage of occupied spermatheca will be higher than the negative control condition. About 20% of the spermathecae were occupied by an oocyte in negative control RNAi while the positive controls (*plc-1* and *gsa-1* RNAi) show more than 95% spermathecal occupancy (Figure 1B).

**Figure 1.**
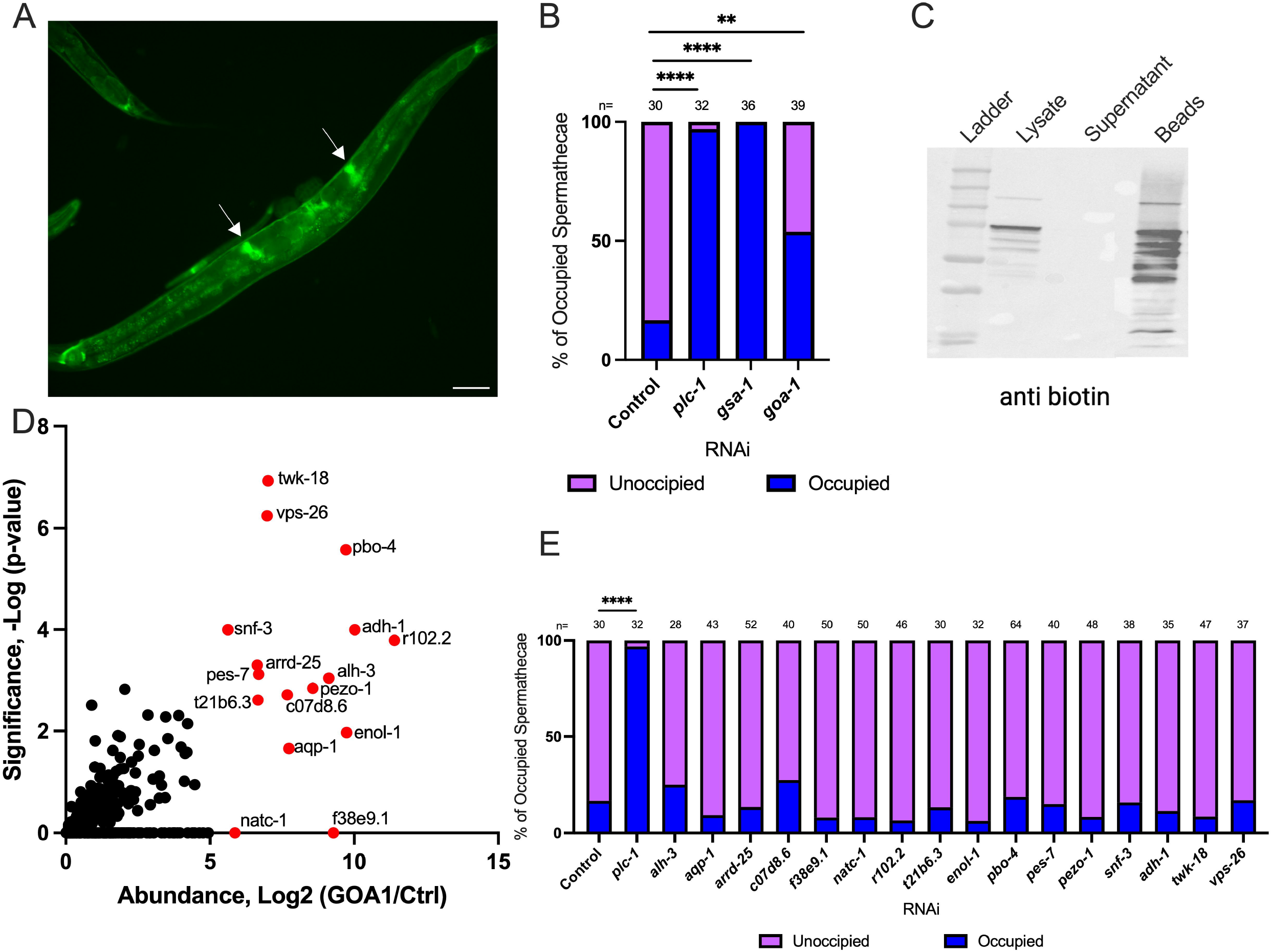
GOA-1 regulates transit of oocytes through the spermatheca. **A)** GOA-1::GFP expression in the hermaphrodite *C. elegans*. Arrows indicate spermathecal expression (scale bar 100 μm). **B)** Spermathecal occupancy assay of *fln-1p*::GCAMP expressing animals grown on negative control (n=30), positive control *plc-1* RNAi (n=32), *gsa-1* RNAi (n=36), and *goa-1* RNAi (n=39). **C)** Western blot detection of biotinylated proteins in the GOA-1::TurboID samples using an anti-biotin antibody. **D)** Right side of a volcano plot indicating significance versus abundance of the detected proteins in three replicate GOA-1::TurboID samples compared to control (*fkh-6p*::TurboID). **E)** Spermathecal occupancy assay of *fln-1p*::GCAMP expressing animals grown on negative control, *plc-1* RNAi, and 16 TurboID targets (n=28-64). Spermathecae were scored for the presence or absence of an embryo (occupied or unoccupied) in the spermatheca. Fisher’s exact t-test (with Benjamini-Hochberg correction) was used for the statistical analysis. Stars designate statistical significance (**** p<0.0001, *** p<0.005, ** p<0.01, * p<0.05).

To identify regulators and effectors of GOA-1 in the spermatheca, potentially including the GPCR, we tagged GOA-1 with the biotin ligase TurboID for proximity labeling (Branon et al., 2018; Cho et al., 2020) under the control of *fkh-6*, a spermathecal-specific promoter (Hope et al., 2003). Western blotting confirmed that GOA-1::TurboID successfully biotinylated proteins (Figure 1C). Mass spectrometry analysis was used to identify the labeled proteins. Animals expressing TurboID alone (*fkh-6p*::TurboID) and wildtype N2 animals were used as controls. We identified 16 proteins that were biotinylated specifically in triplicate GOA-1::TurboID samples. The volcano plot indicates significance versus the abundance of the proteins identified relative to the *fkh-6p*::TurboID control (Figure 1D). The identities and homologies of the TurboID targets are shown in Table 1. We next functionally characterized the GOA-1::TurboID targets for potential roles in transit of oocytes through the spermatheca. We used the same GCaMP expressing strain for each of the 16 genes and scored occupied spermathecae. None of the 16 genes showed a significant spermathecal phenotype (Figure 1E).

**Table 1:**
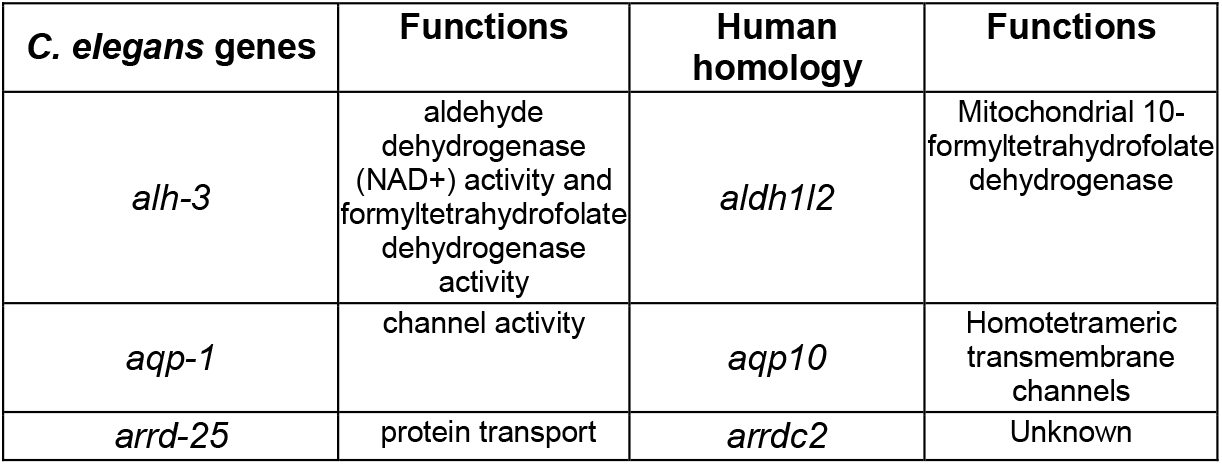
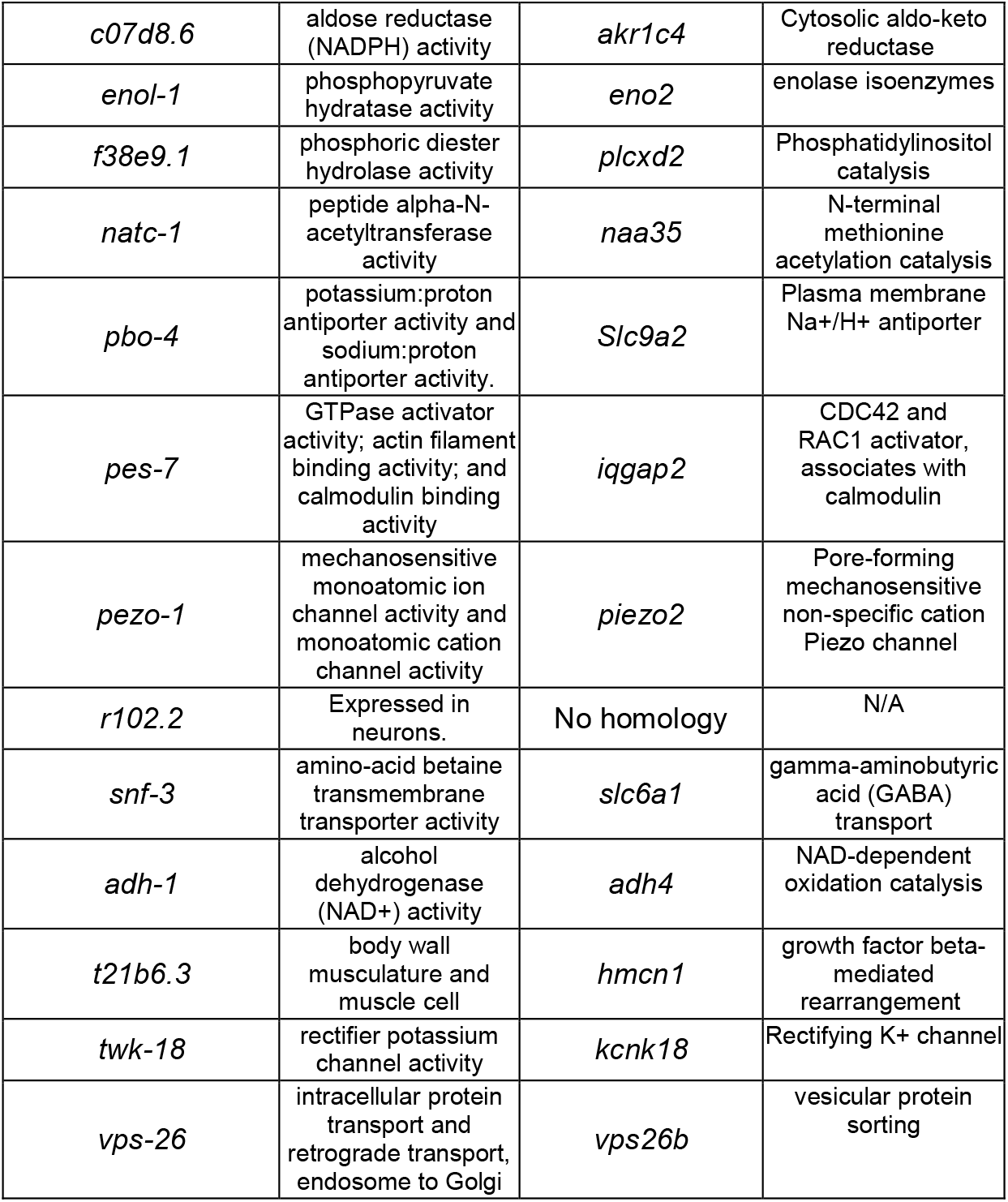
Mass spectrometry results of GOA-1 targets and human homology of the TurboID targets from WormBase and NCBI.

**Table 2:**
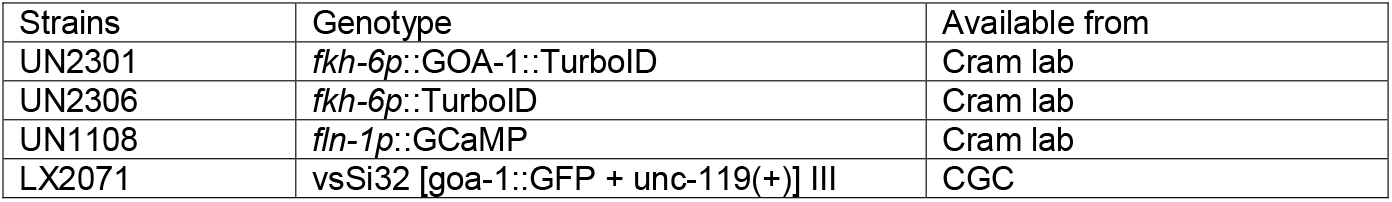
Strains.

**Table 3:**
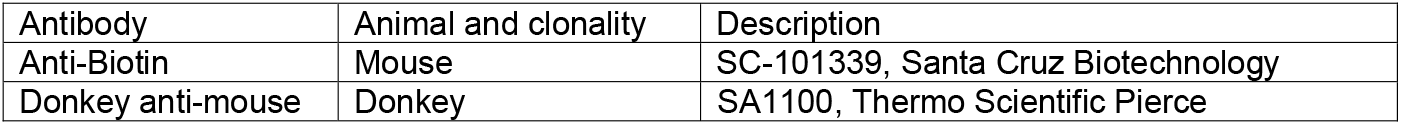
Antibodies.

Although we identified no significant spermathecal phenotype by RNAi, stronger knockdown or analysis of mutant alleles may reveal a role for these genes in the spermatheca. For example, PIEZO1/*pezo-1* does have a known spermathecal phenotype and regulates reproductive tissue contractility (Bai et al., 2020). Some of the other genes may also regulate cell contractility. For example, NATC-1 expression suggests it may be relevant to contractility regulation in the sheath and distal tip cells (Warnhoff et al., 2014). PBO-4 and TWK-18 are expressed in body wall muscle and are expected to regulate muscle contraction (Beg et al., 2008; Kunkel et al., 2000). The next step is to study if these genes show phenotypes with stronger RNAi knock-down or alleles, and if so, to determine whether they interact with GOA-1.

## Methods

### C. elegans strains and culture

Nematodes were grown on NGM plates (0.107 M NaCl, 0.25% wt/vol Peptone, 1.7% wt/vol BD BactoAgar, 2.5 mM KPO_4_, 0.5% Nystatin, 0.1 mM CaCl 2, 0.1 mM MgSO4, 0.5% wt/vol cholesterol) and fed with *E. coli* OP50 at 23°C. All extra-chromosomal arrays of *fkh-6p*::GOA-1::TurboID (20 ng/ul), *fkh-6p*::TurboID (10 ng/ul), and an injection marker (50 ng/ul) were injected into N2 animals. Transgenic animals were integrated by UV radiation. Strains used are *fkh-6p*::GOA-1::TurboID (UN2301), *fkh-6p*::TurboID, (UN2306), *fln-1p*::GCaMP (UN1108), *goa-1*(*sa734*)::GOA-1::GFP (LX2071).

### RNA interference

The RNAi protocol was performed as described previously (Kelley et al., 2020). *HT115(DE3)* bacteria (RNAi bacteria) transformed with a dsRNA construct of interest was grown overnight in Luria Broth (LB) supplemented with 40 μg/ml ampicillin and seeded (150 μl) on NGM plates supplemented with 25 μg/ml carbenicillin and disopropylthio-β-galactoside (IPTG). Seeded plates were left for 24–72 hours at room temperature (RT) to induce dsRNA expression. Empty pPD129.36 vector (“Control RNAi”) was used as a negative control in all RNAi experiments.

Embryos from gravid adults were collected using an alkaline hypochlorite solution as described previously (Castaneda et al., 2020) and washed three times in M9 buffer (22 mM KH_2_PO_4_, 42 mM NaHPO_4_, 86 mM NaCl, and 1 mM MgSO_4_). Clean embryos were transferred to supplemented NGM plates seeded with HT115(DE3) bacteria expressing dsRNA of interest and left to incubate at 23°C for 50–56 hours depending on the experiment.

### Spermathecal occupancy

To prepare age matched young adult animals for the spermathecal occupancy and transit assays, gravid hermaphrodites were lysed in an alkaline hypochlorite solution to release eggs, which were then placed onto seeded NGM plates and grown at 23°C for 52 hours. Animals were mounted on 5% agarose in 0.1 M sodium azide and observed immediately with DIC microscopy to score spermathecal occupancy rates. Imaging was performed on a Nikon Eclipse 80i microscope with a 60× oil-immersion lens using SPOT Advanced software (Version 5.3.5) and a charge-coupled device camera.

### Biotinylation, western blotting, and mass spectrometry

Protein lysates were prepared from age-matched *fkh-6p*::GOA-1::TurboID, *fkh-6p*::TurboID and N2 animals grown on *E. coli HT115(DE3)* as follows: Animals were washed from plates using M9 and allowed to settle and then frozen at -80°C. Animals were resuspended in RIPA lysis buffer containing protease inhibitors (Sanchez & Feldman, 2021). Protein extracts were obtained by sonication at 20% at 10 s intervals for 60 s total. The protein was quantified using the BCA Protein assay. Pierce streptavidin-coated magnetic beads (125ul) were added to 1 mg of protein for each sample (∼250ul), and the lysate was rotated gently at 4°C for 16 h. Then, the samples were washed multiple times with buffers as described (Sanchez & Feldman, 2021).

Samples were loaded onto Mini-PROTEAN TGX precast 10% polyacrylamide gels (Bio-Rad; Hercules, CA, USA) for SDS-PAGE. Antibodies used for Western blotting were Anti-Biotin Antibody at 1:1000 (SC-101339, Santa Cruz Biotechnology) for primary antibody, and donkey anti-mouse at 1:2000 (SA1100, Thermo Scientific Pierce) for secondary antibody.

The eluted samples were sent for proteomic analysis. The proteins were briefly run in a 10% acrylamide gel to clean the samples followed by in gel reduction, alkylation and trypsin digestion (Shevchenko et al, Nature Protocols, 2006; 1(6):2856). The resulting peptides were desalted with a C18 column (Pierce), dried down and reconstituted with 0.1% formic acid. Proteomic analysis was carried on a ThermoFisher QExactive Orbitrap in line with a Thermo Fisher RSLC Ultimate 3000 nanoUPLC. The peptides were loaded onto an in-house pulled tip 75 μm x 20 cm C18 ReproSil-Pur 120 1.9 μm (Dr. Maisch) LC column and separated with the following gradient: buffer A; 0.1% formic acid, buffer B; acetonitrile with 0.1% formic acid, 0-25 min: 2% B, 25-14 5min: 2-35% B, 145-154min: 35-95% B, 154-159min: 95% B, 159-160 min: 95-2% B, 160-180 min: 2% B. The flow rate was set to 200 nL/min. The QExactive settings were as follow, spray voltage 1.5 kV, sheath gas flow rate 10, capillary temperature 250°C, S-lens RF level 65. The data was acquired using a top 20 data dependent acquisition scan with the following parameters: for MS1, the resolution was set at 70,000, AGC target at 3e6, max IT 100 ms with a scan range from 350 to 1500 m/z. For MS2, the resolution was set at 17,500, the AGC target at 1e5 and the maximum IT at 100 ms with an isolation window of 1.6 m/z and a collision energy of 30. The resulting spectra were analyzed with proteome discoverer 3.0 with a canonical *C. elegans* FASTA file containing 4462 proteins, a common contaminant database, cysteine carbamidomethyl (+57.02 Da) set as a fixed modification, methionine oxidation (+15.99 Da) set as a dynamic modification and acetylation (+42.01 Da) and methionine loss (-131.04 Da) as protein terminus dynamic modifications. Proteins were filtered to 1% false discovery rate. A list of proteins identified by mass spectrometry in three replicates is shown in Table 1.

### Image analysis and Statistics

ImageJ (FIJI) (version 2.16.0) was used for image analysis. GraphPad Prism (version 10.5.0) was used for statistical analyses. Phenotype distributions were evaluated with Fisher’s exact test. In both cases, The Benjamini-Hochberg correction was applied to adjust for multiple testing.

